# A Guided AI Framework for Customizable and Efficient Harmonization to the OMOP Common Data Model

**DOI:** 10.64898/2026.08.07.742453

**Authors:** Nishu Nehra, Rohit Swami, Dharani Dadi, Ritika Mishra, Umang Sharma, Pawan Verma, Manimala Sen, Nobal Kishor, Abhishek Jha

## Abstract

Getting clinical data from different sources to “talk” to each other within the OMOP Common Data Model (CDM) is arguably the most tedious part of multi-center research. While this integration is essential, the transformation process is frequently a manual grind, requiring a rare overlap of deep clinical knowledge and technical expertise. In this paper, we present a framework designed to alleviate some of the burden on the researcher by automating data harmonization through two distinct steps: structural schema mapping and terminological standardization.

For the structural piece, we moved away from “black box” logic in favor of a stateful workflow managed by large language models (LLMs) and directed acyclic graphs. By profiling EHR data at the source, our system generates context-aware dictionaries that offer ranked mapping suggestions alongside confidence scores. While our benchmarking showed a 97.5% agreement rate at the schema level and an 84% agreement rate at the value level when compared with human experts, the system appears most effective when treated as a “co-pilot” rather than a total replacement for human oversight.

To handle value-level standardization, we implemented a hybrid search strategy that pairs the semantic depth of SapBERT embeddings with the literal precision of fuzzy string matching. By using FAISS for rapid similarity retrieval, the engine attempts to resolve messy or “noisy” clinical descriptions to standard OMOP concepts. This approach seems particularly promising for handling the non-standardized labels that often plague smaller, local datasets.

Ultimately, our results suggest that this guided approach can shift the timeline for OHDSI-compliant warehousing from weeks of manual curation to a more manageable and scalable pipeline, potentially lowering the barrier to entry for smaller research teams.

## 1 Introduction

For over a decade, the promise of large-scale observational research has hinged on our collective ability to speak a common data language, specifically, the OMOP Common Data Model [1]. By standardizing disparate Electronic Health Record (EHR) systems, the OHDSI community has enabled studies that span continents and millions of patients. Yet, those who have spent time in the “trenches” of data curation know that the initial Extract, Transform, Load (ETL) process remains a hurdle. Despite the availability of tools like White Rabbit [2], Rabbit-in-a-Hat [2], and Usagi [3], the reality of data integration is still largely a manual, laborious process that demands a rare intersection of clinical nuance and technical expertise [4].

The core of the problem isn’t just the sheer volume of data, but its inherent messiness. Clinical records are often a chaotic mix of local terminologies, idiosyncratic abbreviations, and unstructured notes that resist simple rule-based mapping. While the recent surge in Large Language Models (LLMs) has offered a tempting alternative for medical data processing [5], their “black box” nature can be a significant liability in a research context. We cannot simply trust an unverified model to decide how a local lab result should be represented in a global study. There is a persistent and arguably justified skepticism regarding the transparency and reproducibility of AI-driven harmonization.

This paper grew out of a need to balance that AI-driven efficiency with the “human-in-the-loop” oversight that clinical research demands. We propose a framework that moves away from monolithic, one-shot automation in favor of a tiered approach. At the schema level, we utilize a stateful workflow that attempts to mimic the reasoning of a domain expert, generating transparent, ranked recommendations rather than opaque “final” answers. For terminological mapping, we found that combining the semantic depth of biomedical embeddings generated by SapBERT [6] with traditional fuzzy matching helps capture nuances that either method alone might miss.

Our goal was not to build a system that replaces the human expert, but rather one that handles the “heavy lifting” of data profiling and similarity search, allowing the researcher to focus on the high-level validation that ensures data integrity. In the following sections, we describe how this guided, parallelized approach might help turn a months-long curation ordeal into a more manageable and scalable pipeline.

## 2 Methods

### 2.1 System Architecture and Workflow Orchestration

The system is implemented as a cloud-native, event-driven pipeline designed to support the scale, heterogeneity, and privacy constraints of real-world electronic health record (EHR) data. At a high level, the architecture separates the workflow into a presentation layer for project management and expert review, a processing layer for asynchronous data transformation and mapping, and a storage layer for both raw and derived artifacts. When a user uploads one or more clinical source tables, the files are registered within a project-scoped workspace and stored as immutable source assets, while associated metadata are recorded to preserve provenance and support downstream orchestration. This design enables the platform to manage multi-file studies in a reproducible manner while maintaining clear separation between raw inputs, intermediate artifacts, and finalized outputs.

Rather than processing each dataset as a single monolithic task, the workflow is executed as a stateful directed acyclic graph (DAG) (Figure 1) in which each stage produces explicit outputs that are consumed by subsequent stages. This orchestration model decomposes the pipeline into discrete, auditable steps, including source ingestion, asset preparation, candidate OMOP [1] mapping generation, expert curation, vocabulary normalization, and preview materialization. The stateful nature of the DAG is important because it allows the system to preserve intermediate decisions and artifacts across stages, including file-level profiling results, governance annotations, candidate mapping sets, and finalized review outcomes. As a result, each stage can be executed, monitored, and, when necessary, repeated independently without requiring the entire workflow to be rerun from the beginning.

**Figure 1:**
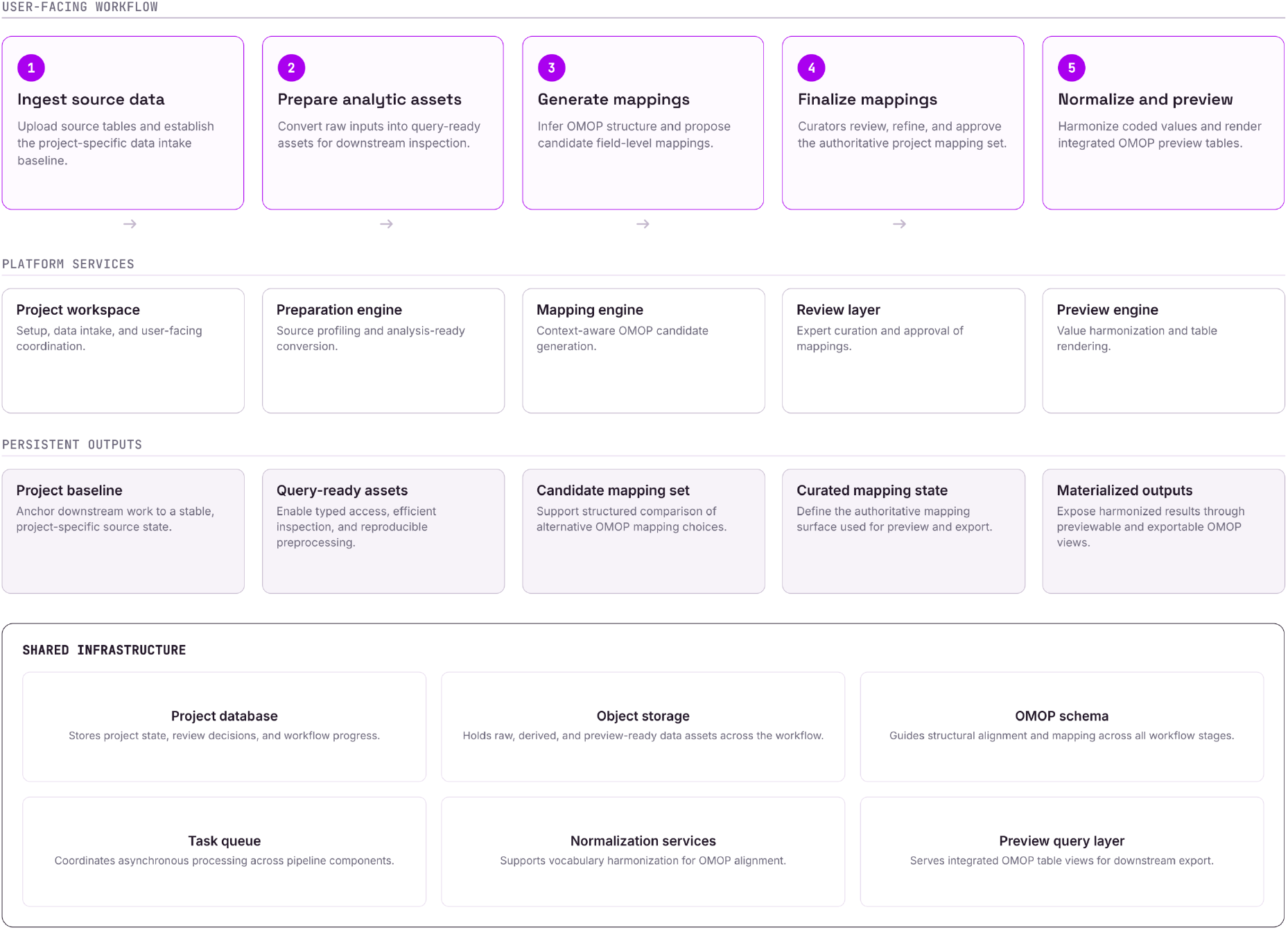
Data onboarding and mapping pipeline. Five user-facing stages (ingest, prepare, generate, finalize, normalize) map to their supporting platform services, persistent outputs, and shared infrastructure components.

#### 2.1.1 Source Data Ingestion

The workflow starts by importing source tables into a project-level workspace. These files may originate from different operational systems and often differ in naming conventions, structure, and completeness. During ingestion, each file is registered as a project asset so that it can be tracked throughout the workflow. This step establishes the raw input layer for downstream profiling, governance, mapping, and review.

#### 2.1.2 Data Profiling

After ingestion, the platform transforms the raw files into analysis-ready assets and captures their structural metadata. This includes identifying columns, preserving file-level organization, and preparing the data in a format that supports efficient inspection and query. The purpose of this step is to standardize heterogeneous source tables into a more consistent computational form, making later stages such as schema alignment, preview generation, and mapping more robust and scalable.

#### 2.1.3 Mapping Generation using LLM

For non-sensitive columns, the platform generates candidate mappings to OMOP [1] tables and fields. This step uses the structure and content of the source data, together with project-level context across multiple files, to infer the most likely target locations in the OMOP [1] model. Rather than producing a single fixed answer, the system creates a ranked set of plausible mappings, which allows uncertainty to be preserved and reviewed explicitly. This is especially important when multiple source columns could map to related OMOP [1] domains or when source semantics are only partially specified.

#### 2.1.4 Vocabulary normalization and OMOP preview generation

After final mapping decisions were established, the workflow materialized an OMOP-aligned representation of the source data. For fields requiring terminology harmonization, source values were normalized into standardized concept-level representations so that semantically equivalent entries could be encoded consistently. The finalized mappings were then applied to generate an integrated preview of the projected OMOP [1] tables, allowing users to inspect the transformed data before export or downstream analysis. This final stage served both as a harmonization step and as a validation layer, making it possible to assess how the curated mapping logic behaved when executed on the original source records.

### 2.2 The Hybrid Concept Mapping Engine

This component of the pipeline addresses the normalization of local source terms to OMOP [1] Standard Concepts, which is a central requirement for semantic interoperability within the OMOP Common Data Model [1]. Because local EHR terms are often incomplete, abbreviated, institution-specific, or lexically inconsistent, concept normalization cannot rely on exact string matching alone. The proposed engine (Figure 2) therefore uses a hybrid retrieval strategy that combines biomedical semantic embeddings with lexical similarity scoring, followed by OMOP-aware filtering and reranking to improve mapping precision and standards compliance.

**Figure 2:**
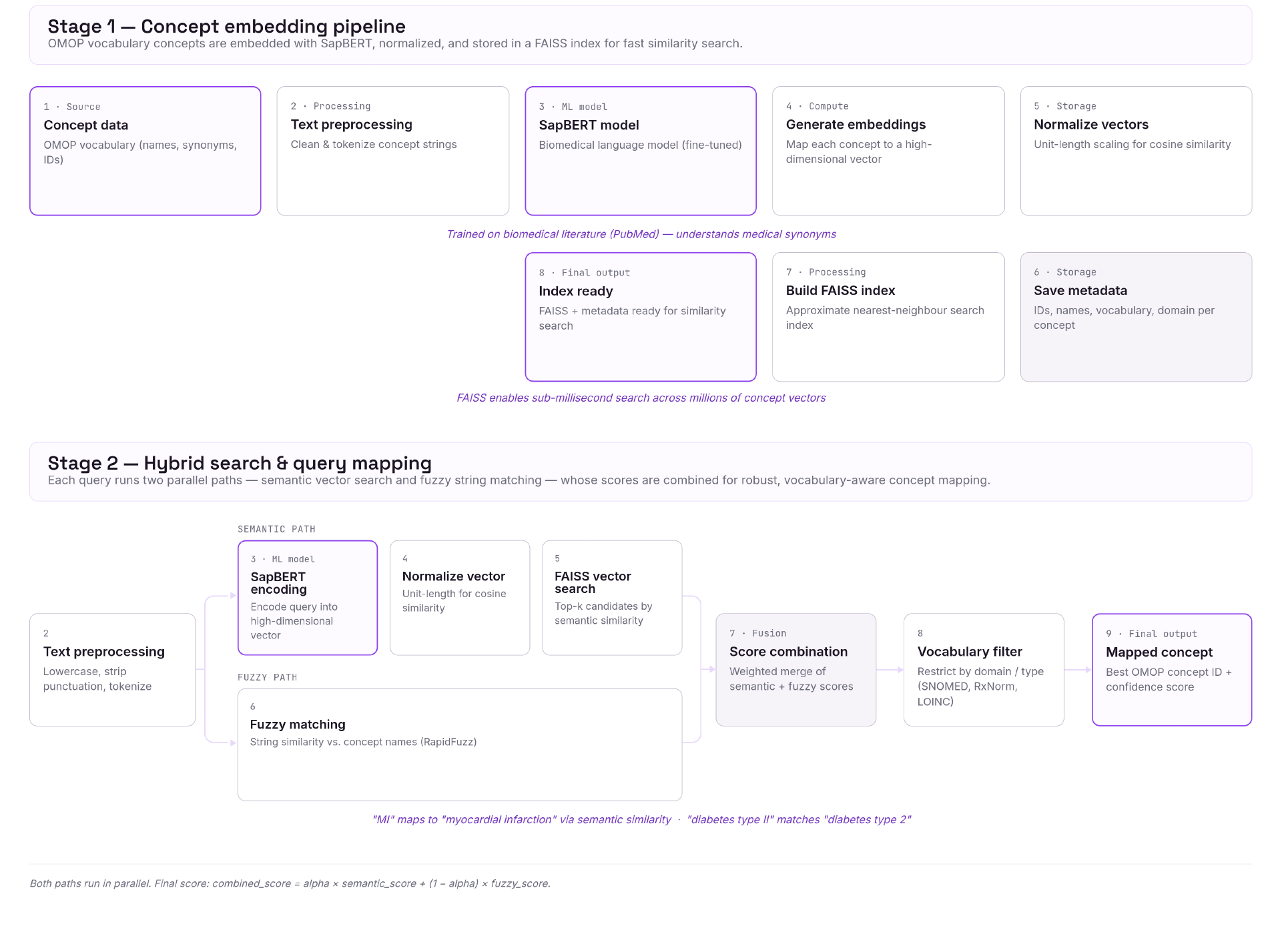
OMOP concept mapping via hybrid search. Stage 1 builds a SapBERT-embedded FAISS index over OMOP vocabulary concepts; Stage 2 maps incoming query terms by combining semantic vector search with fuzzy string matching into a single confidence-scored concept.

#### 2.2.1 OMOP Vocabulary as the Retrieval Substrate

The mapping engine is grounded in the OMOP [1] standardized vocabulary, in which clinical meaning is represented through concepts, synonyms, and explicit mappings between non-standard source concepts and standard target concepts. In OMOP [1], Standard Concepts are the concepts used to populate normalized CDM fields, whereas non-standard concepts are retained as source concepts and linked to standard targets through vocabulary relationships such as Maps to and Maps to value. This vocabulary structure makes concept normalization a constrained retrieval problem: the objective is not merely to find a textually similar term, but to recover the most appropriate standard clinical concept within the OMOP [1] framework.

#### 2.2.2 Embedding Generation and Index Construction

To support semantic retrieval, OMOP [1] concept strings were transformed into dense vector representations using SapBERT [6], a biomedical representation model specifically developed to improve entity-level synonym alignment in the clinical and biomedical domain. SapBERT [6] is particularly suitable for this task because it was designed to place synonymous biomedical entities closer in the representation space. This improved the performance of the medical entity linking process. These concept embeddings were then normalized and indexed for efficient large-scale similarity search. For vector retrieval, the system uses FAISS [7], which provides efficient similarity search over dense vector collections and supports scalable nearest-neighbor retrieval over high-dimensional embeddings.

#### 2.2.3 Dual-Path Query Processing

At inference time, each source term is processed through two complementary retrieval pathways. In the semantic pathway, the query term is encoded using the same SapBERT [6] model, normalized in the embedding space, and submitted to the vector index to retrieve a candidate set on the basis of semantic similarity. This branch is intended to capture contextual and conceptual equivalence, including abbreviations, synonyms, and clinically related expressions that may not closely match at the surface-text level. In parallel, the lexical pathway computes a fuzzy string similarity score between the query and candidate concept names. This branch preserves literal textual evidence and improves robustness to spelling variation, token reordering, and near-exact string matches. The use of these two branches reflects the observation that semantic similarity and lexical similarity solve different failure modes in concept normalization, and that combining them provides a more reliable retrieval signal than either method alone.

#### 2.2.4 Score Fusion and Candidate Ranking

The final candidate ranking is obtained by combining the semantic and lexical signals into a unified score. Let *s*_emb_(*q, c*) denote the embedding-based similarity between query *q* and candidate concept *c*, and let *s*_fuzzy_(*q, c*) denote the lexical similarity score. The combined score is defined as:

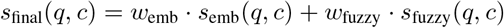

where *w*_emb_ and *w*_fuzzy_ are tunable coefficients controlling the relative contribution of semantic and lexical evidence. In the current implementation, the semantic branch is given higher weight than the fuzzy branch, reflecting the need to recover conceptually correct matches even when local source terms are abbreviated or institution-specific, while still retaining textual evidence as a safeguard against semantically plausible but lexically unsupported results. This weighted fusion provides a practical balance between semantic generalization and lexical fidelity.

#### 2.2.5 OMOP-Aware Filtering and Final Concept Selection

After score fusion, the ranked candidate list is further constrained using OMOP-specific metadata [1]. Candidates may be filtered by target vocabulary, allowing normalization to be restricted to the intended annotation library, such as SNOMED [8], RxNorm [9], or LOINC [10]. In addition, Standard Concepts are prioritized over non-standard concepts so that the selected mapping is directly compatible with OMOP-normalized data representation [1]. This post-retrieval filtering step is important because it ensures that the final output is not only semantically similar to the source term, but also operationally valid within the OMOP [1] vocabulary system. In this way, the engine functions as a vocabulary-aware hybrid retrieval model rather than a generic clinical text similarity tool.

#### 2.2.6 Evaluation Metrics

The evaluation was conducted separately for the two core normalization tasks addressed by the pipeline: schema mapping and terminology harmonization. For schema mapping, the predicted alignments between source fields and target OMOP [1] tables or columns were compared against the reference mappings derived from the ground-truth dataset. For terminology harmonization, the predicted standardized concepts were assessed against the corresponding reference concepts in the same dataset. This two-level evaluation allowed us to distinguish structural correctness at the schema level from semantic correctness at the vocabulary level.

To assess the reproducibility of the LLM-assisted components, the full pipeline was executed five independent times under the same experimental conditions for both schema mapping and terminology harmonization. Because large language model outputs can exhibit residual stochasticity even under controlled settings, repeated execution was used to evaluate run-to-run stability rather than assuming strict determinism. Agreement across runs was therefore used as an indicator of output consistency and robustness.

All predicted mappings were benchmarked against the Synthea10k COVID-19 [11] reference dataset, which contains synthetic EHR records for approximately 10,000 patients with COVID-19 or related clinical conditions. This dataset was used as the evaluation ground truth because it provides a controlled setting with known source-to-target correspondences, enabling systematic assessment of both field-level schema alignment and concept-level terminology normalization.

### 2.3 Support for Custom Data Models and Multi-Modal Harmonization

Although the primary implementation targets OMOP CDM [1], the architecture was designed as a schema-driven harmonization framework rather than as an OMOP-only mapper (Figure 3). This design choice allows the same orchestration and mapping logic to be applied to alternative target data models, provided that the target schema can be expressed in a machine-readable form. In addition, the project-level workflow is able to integrate heterogeneous source tables within a single harmonization task, enabling coordinated mapping across multiple clinical data modalities represented in structured tabular form.

**Figure 3:**
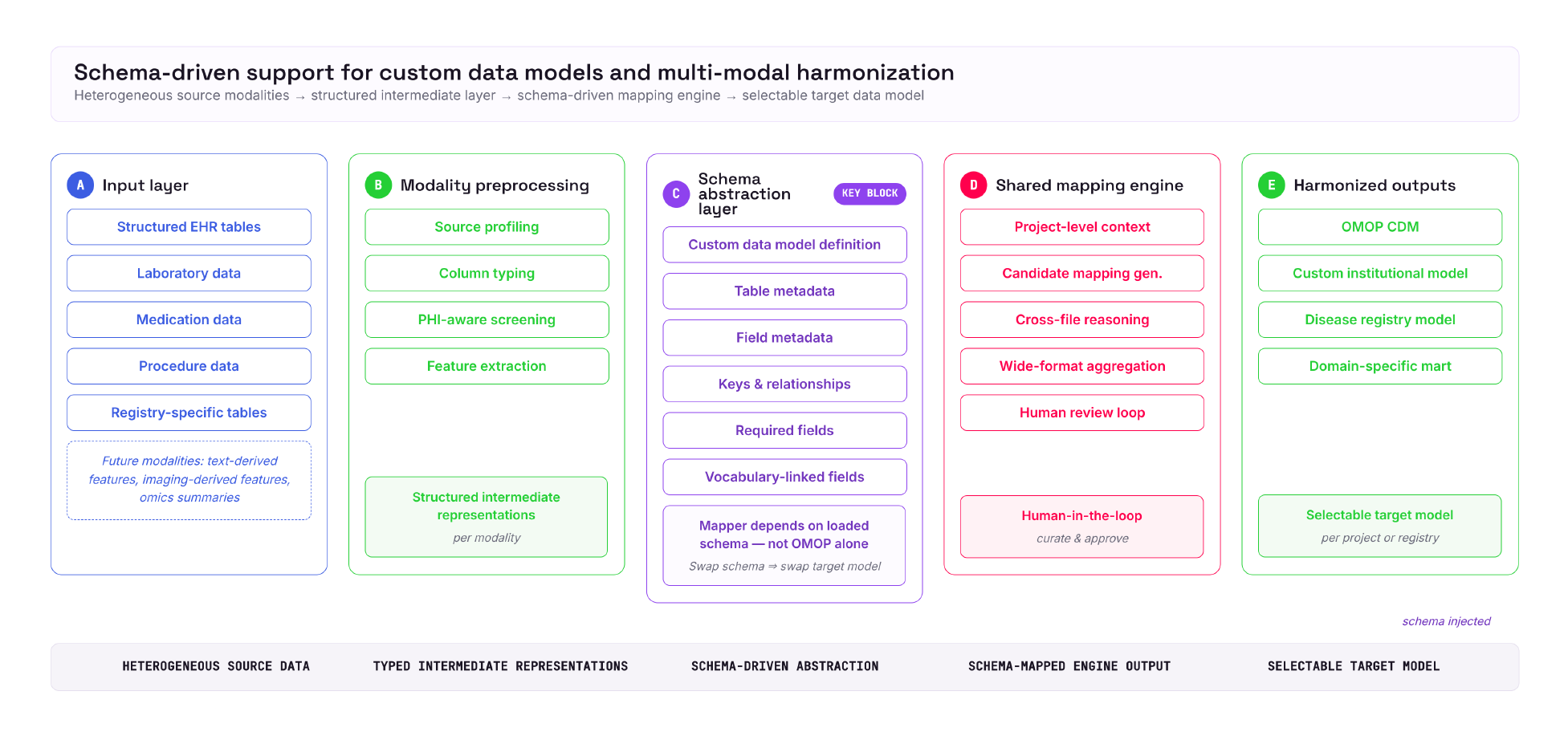
Schema-driven harmonization across custom data models. Heterogeneous source modalities pass through preprocessing into a schema abstraction layer (block C), which drives a shared mapping engine to produce a harmonized output in any selectable target data model, not OMOP alone.

#### 2.3.1 Abstraction of Target Model Semantics

Support for custom data models is achieved by externalizing the target schema from the core mapping logic. Instead of hardcoding table names, field names, or OMOP-specific assumptions throughout the pipeline, the system loads model definitions from explicit schema metadata that describe tables, columns, data types, required fields, primary keys, foreign-key relationships, and concept-linked fields. As a result, the mapper operates over a generalized representation of the target model and queries schema metadata dynamically during candidate generation, conflict checking, and preview construction. This allows the same pipeline to be re-targeted to alternative harmonization frameworks, including institution-specific analytical models, research registries, domain-focused clinical marts, or other standardized healthcare schemas, with minimal changes to the orchestration layer itself.

From a methodological perspective, this abstraction is important because it separates mapping strategy from target schema definition. The target model becomes a configurable input to the system rather than a fixed implementation constraint. In practice, adapting the workflow to a custom model requires defining the schema in the expected structured format and specifying the relevant field-level metadata that guide mapping and validation. Once that schema is available, the same project-level processes for source profiling, candidate mapping, review, and output materialization can be reused. This design makes the framework suitable not only for OMOP-based standardization, but also for settings in which local institutions require harmonization into custom downstream representations.

#### 2.3.2 Harmonization Across Heterogeneous Clinical Modalities

The platform also supports harmonization across multiple structured clinical data modalities within the same project. Here, “multi-modal” refers to the integration of heterogeneous but related data sources, such as demographics, diagnoses, medications, laboratory measurements, procedures, observations, and other tabular clinical domains that may originate from different source systems or files. Rather than treating each file independently, the workflow constructs project-level context across all uploaded sources and uses that shared context during mapping generation. This enables the system to infer how related variables from separate tables contribute to a unified target representation.

#### 2.3.3 Cross-File Contextual Mapping and Aggregation

This capability is particularly relevant for real-world EHR harmonization, where clinically meaningful information is often distributed across multiple relational tables or exported files rather than stored in a single source dataset. The pipeline therefore supports cross-file reasoning at the project level and can preserve many-to-one mapping patterns when multiple source columns contribute to the same target variable. Such behavior is useful for wide-format or fragmented source representations, where semantically related attributes may need to be aggregated, normalized, or transformed before they can be expressed in the target model. In this sense, the framework is not limited to one-table-to-one-table translation, but can support more complex harmonization patterns across heterogeneous structured inputs.

#### 2.3.4 Extensibility beyond current structured modalities

While the present implementation is optimized for structured tabular EHR data, the same schema-driven design provides a foundation for broader multi-modal harmonization. Additional modalities, such as clinical text, imaging-derived variables, waveform summaries, or genomics-derived features, could be incorporated by introducing modality-specific pre-processing steps that transform those inputs into structured intermediate representations compatible with the target schema. Under this formulation, the harmonization engine remains unchanged at its core, while modality-specific encoders or extractors serve as upstream components that generate schema-aligned features for downstream mapping and normalization. This makes the overall framework extensible to richer clinical data environments without requiring a complete redesign of the mapping architecture.

## 3 Results and Discussion

The present work was designed not only as an automated mapping pipeline, but as an interactive harmonization workflow in which computational inference and curator oversight are tightly coupled. Rather than treating standardization as a single transformation step, the workflow separates data understanding, schema alignment, terminology harmonization, and output validation into distinct but connected phases. This is important because real-world clinical datasets are rarely difficult for only one reason. In practice, ambiguity can arise from inconsistent source schemas, incomplete metadata, local coding practices, and differences in how similar clinical events are recorded across systems. By organizing these challenges into a guided workflow, the platform supports a more interpretable and auditable path from heterogeneous source tables to OMOP-aligned data [1].

### 3.1 Data Profiling

The first important contribution of the workflow is the data profiling stage (Figure 4), which helps establish context before any mapping decision is made. Profiling provides a structured summary of the uploaded tables, including field composition, sparsity, uniqueness patterns, and representative values. From a harmonization perspective, this stage is more than a descriptive convenience: it functions as the interpretive layer that helps users distinguish clinically meaningful variables from administrative, redundant, or poorly populated fields. This improves downstream mapping quality because candidate alignments are informed not only by field names, but also by the statistical and semantic characteristics of the source data. In this sense, data profiling reduces the risk of premature or weakly justified mappings and helps ground later automated decisions in observable source evidence.

**Figure 4:**
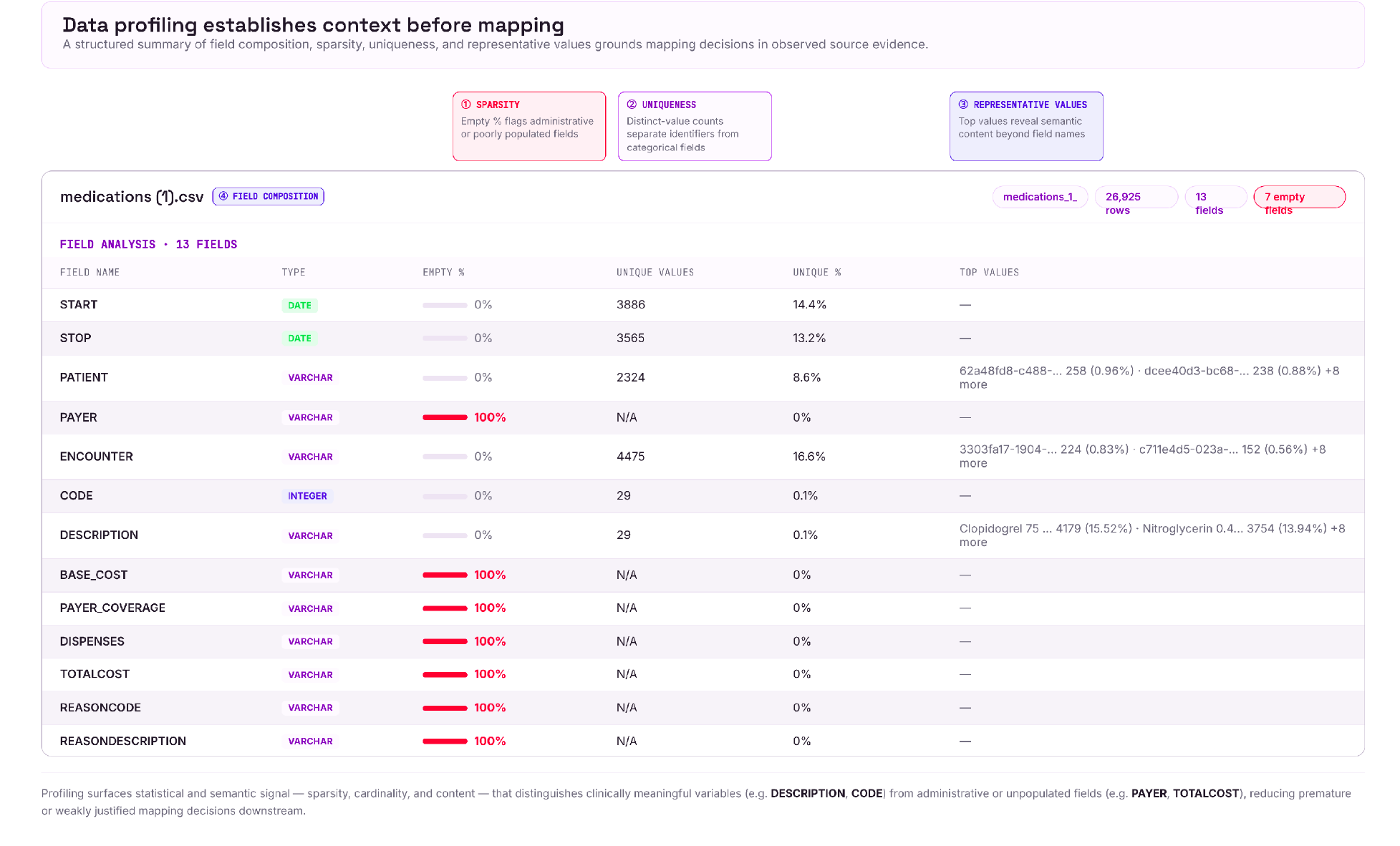
Data profiling interface used during source-table review. The view summarizes uploaded tables through field-level metadata, representative values, sparsity and distinctness cues, and table-specific context that helps users identify informative variables before schema mapping and terminology harmonization.

### 3.2 Automated Mapping and Scoring

The schema-mapping stage (Figure 5) illustrates the practical value of combining automation with explicit review. Candidate mappings are generated as ranked suggestions rather than as irreversible assignments, which allows the system to narrow the decision space while preserving uncertainty for curator inspection. This is especially useful in EHR harmonization, where one source field may plausibly correspond to several related target fields depending on provenance, granularity, or intended analytic use. Score-guided ranking provides a transparent basis for review and helps users prioritize high-confidence alignments while examining ambiguous cases more carefully. As a result, automation improves efficiency without removing the role of expert judgment, which remains essential for resolving clinically nuanced or institution-specific fields.

**Figure 5:**
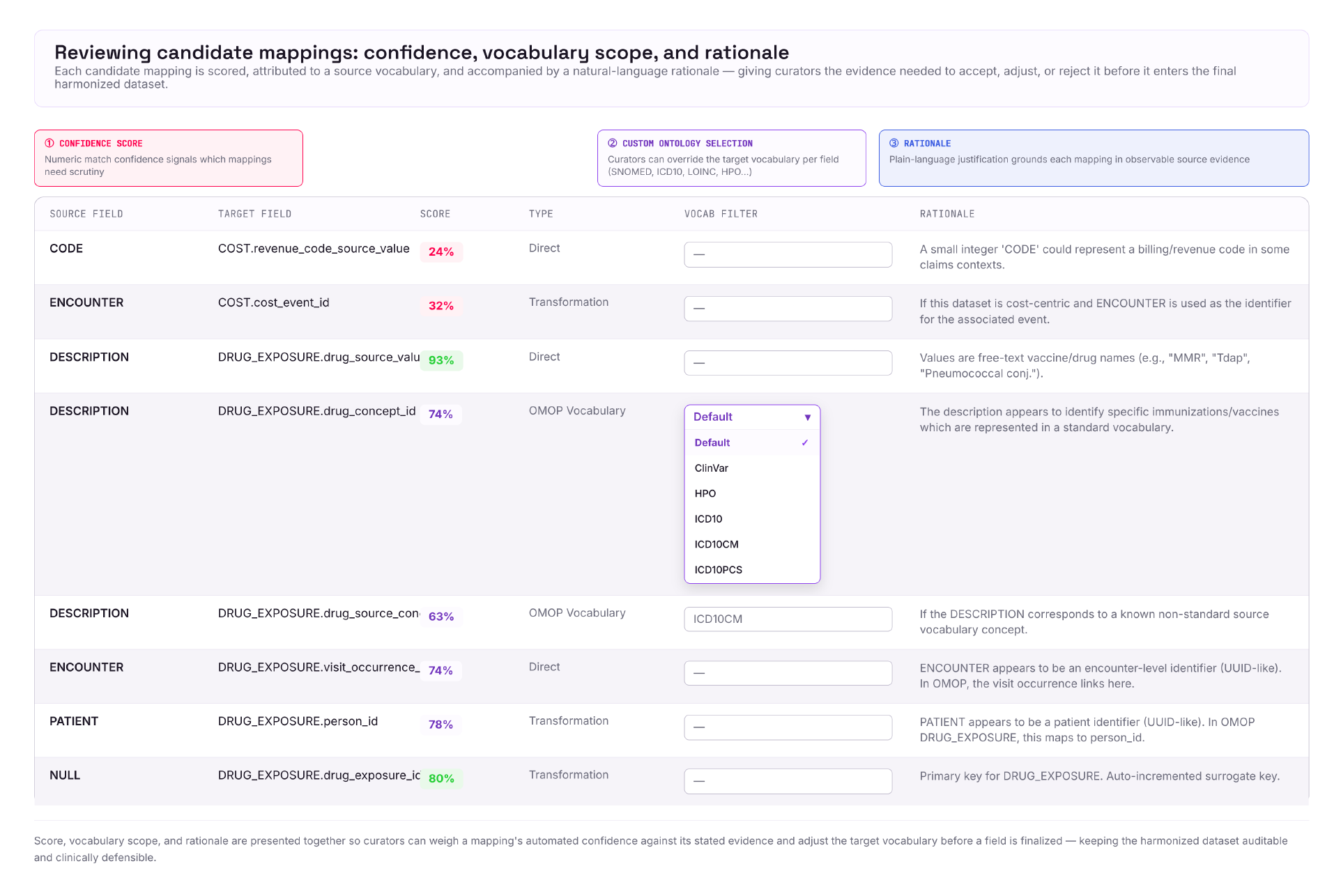
Mapping review interface for ontology-constrained harmonization. The workspace presents candidate concept matches with scoring evidence, source-term context, and reviewer controls so users can validate or revise value-level mappings before finalizing normalization for the selected field.

### 3.3 Custom Selection of Ontologies for Harmonization of Relevant Fields

A second important feature of the workflow is the use of field-aware terminology harmonization (Figure 5). In practice, clinical value normalization is most reliable when candidate concepts are interpreted in the context of the destination field rather than through unconstrained concept search alone. This allows harmonization to remain aligned with the semantic role of the target variable, whether the relevant concept space is demographic, diagnostic, procedural, laboratory, or medication-related. Such field-constrained normalization is methodologically valuable because it reduces the likelihood of semantically related but operationally inappropriate matches. More broadly, it supports the view that terminology harmonization should be guided by schema context and intended use, rather than treated as an isolated vocabulary lookup problem.

### 3.4 Benchmarking Using Synthea COVID-19 10K Data

Benchmarking on the Synthea COVID-19 10K dataset [11] provided a useful assessment of the framework at both the schema and value levels. The evaluation was performed over five independent runs to assess whether the system produced consistent mappings at each stage.

At the schema level (Table 1), the results indicate strong recovery of target structure across the evaluated OMOP tables [1], supporting the system’s ability to align heterogeneous source tables with the expected analytical representation. At the value level (Table 2), performance remained favorable across the clinically informative and directly comparable domains included in the evaluation. The benchmark showed consistent concept-level harmonization across key domains such as condition occurrence, drug exposure, person, procedure occurrence, and provider. Device exposure was excluded from interpretation, and visit occurrence was not included in the value-level summary because it did not provide a meaningful basis for direct concept-level comparison in this evaluation setting.

**Table 1:** Evaluation of Schema-Level Harmonization of Synthea COVID19 tables against Synthea COVID19 ground truth.

| OMOP CDM Table | Raw Source Table | Mapping Accuracy (%) |
| --- | --- | --- |
| Condition Occurrence | Conditions | 100.00 |
| Device Exposure | Devices | 78.95 |
| Drug Exposure | Medications, Immunizations | 100.00 |
| Person | patients | 100.00 |
| Procedure Occurrence | procedures | 100.00 |
| Provider | providers | 100.00 |
| Visit Occurrence | encounters | 61.90 |
| **Mean Accuracy** |  | **97.50** |

**Table 2:** Evaluation of Value-Level Harmonization of Synthea COVID19 tables against Synthea COVID19 ground truth.

| OMOP Table | Raw Source Table | Raw Source Field | Mapping Accuracy (%) |
| --- | --- | --- | --- |
| Condition Occurrence | Conditions | Code | 94.60 |
| Drug Exposure | Medications, Immunizations | Code | 81.98 |
| Person | Patients | Race, Ethnicity, Gender | 99.10 |
| Procedure Occurrence | Procedures | Code | 80.00 |
| Provider | Providers | Gender, Specialty | 63.80 |
| **Mean Accuracy** |  |  | **83.89** |

**Table 3:** List of figures and tables.

| Figure/Table | Title |
| --- | --- |
| Figure 1 | Data onboarding and mapping pipeline |
| Figure 2 | OMOP concept mapping via hybrid search |
| Figure 3 | Schema-driven support for custom data models and multi-modal harmonization |
| Figure 4 | Data profiling interface for source-table inspection and field-level evidence |
| Figure 5 | Mapping review interface for ontology-constrained harmonization |
| Table 1 | Schema-level accuracy by OMOP table |
| Table 2 | Value-level accuracy by OMOP table |

**Table 4:** List of abbreviations.

| Abbreviation | Full Form |
| --- | --- |
| CDM | Common Data Model |
| COVID-19 | Coronavirus Disease of 2019 |
| DAG | Directed Acyclic Graph |
| EHR | Electronic Health Record |
| ETL | Extract, Transform, Load |
| FAISS | Facebook AI Similarity Search |
| LLM | Large Language Model |
| LOINC | Logical Observation Identifiers Names and Codes |
| OHDSI | Observational Health Data Sciences and Informatics |
| OMOP | Observational Medical Outcomes Partnership |
| SNOMED | Systematized Nomenclature of Medicine |

At the schema level (Table 1), the benchmarking results indicate that the pipeline achieved strong and largely stable structural alignment between the source data and the target OMOP schema across repeated runs. Core domains such as person, condition, drug, and provider were mapped with consistently high fidelity, suggesting that the workflow is able to preserve the intended target-table assignment once the source structure is profiled and the candidate ontologies are selected. The remaining variation at schema level (Appendix Figure 6) appears to be driven less by instability in the pipeline and more by inherent differences between source-table semantics and OMOP’s domain boundaries, particularly in domains where a single source construct may plausibly map to more than one standardized representation. Overall, the table-level results support the reliability of the UI-guided harmonization journey for establishing reproducible schema alignment with limited manual intervention.

**Figure 6:**
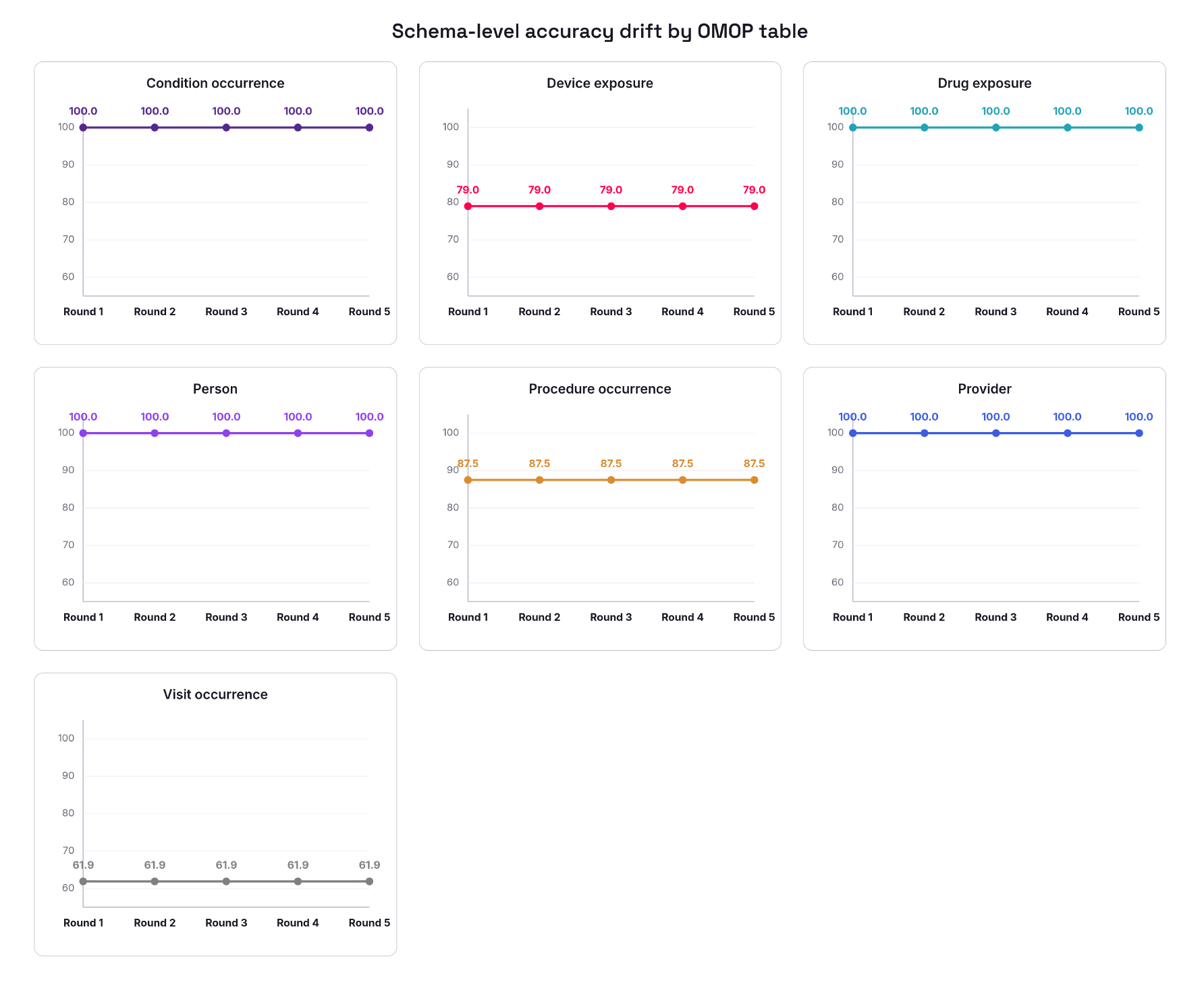
Evaluation of Schema-Level Harmonization of Synthea COVID19 tables against Synthea COVID19 ground truth with per-run accuracy. Most tables hold stable at or near 100%; Device Exposure, Procedure Occurrence, and Visit Occurrence plateau at lower, table-specific ceilings (79%, 87.5%, and 61.9% respectively).

At the value level (Table 2), the results show that concept harmonization remained strong overall, with especially robust performance in domains where source values are relatively well-constrained and semantically explicit, and with clear improvement across iterative runs in medication-related mappings as the workflow better resolved ontology choices and source-term normalization. The observed drift across runs (Appendix Figure 7) was modest rather than systemic, indicating that most fluctuations arose from edge-case terminology, concept granularity differences, and vocabulary-specific naming conventions rather than from failure of the mapping framework itself. These findings suggest that the main areas for improvement are targeted rather than foundational: strengthening context-aware concept disambiguation, improving normalization of source terms before ontology lookup, and adding domain-specific validation checks that can flag near-match concept selections before final harmonization. Together, these refinements would further improve consistency while preserving the strong overall accuracy already demonstrated by the system.

**Figure 7:**
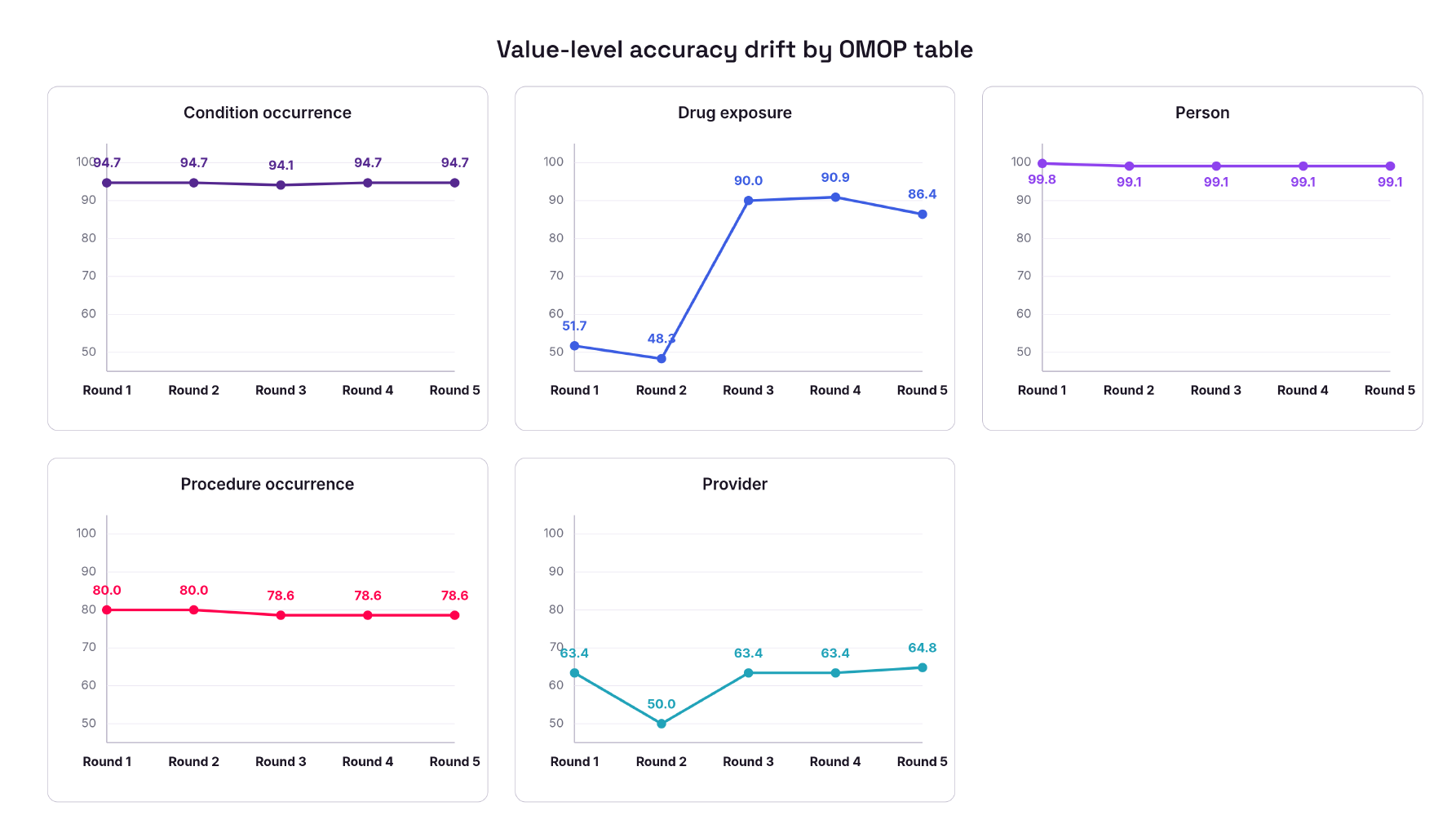
Evaluation of Value-Level Harmonization of Synthea COVID19 tables against Synthea COVID19 ground truth with per-run accuracy. Person and Condition Occurrence remain consistently high (>94%); Drug Exposure improves sharply after Round 2; Provider accuracy stays comparatively low and variable.

Taken together, these findings support a positive interpretation of the framework’s harmonization performance. The system performed well not only in assigning source fields to the appropriate structural destinations, but also in preserving clinically meaningful semantic alignment across the major value-level domains that were evaluated. This is important for practical use, where the utility of harmonization depends on both structural correctness and the reliability of standardized values used in downstream analysis. Overall, the benchmark suggests that the framework is capable of producing analytically useful standardized outputs across both schema-level and value-level evaluation settings.

## 4 Conclusion

This study presents a human-in-the-loop framework for EHR harmonization that integrates data profiling, automated schema mapping, field-aware terminology normalization, and OMOP-oriented output validation within a single workflow. The overall design treats harmonization as a guided sequence of semantic and structural decisions, allowing automation to accelerate routine alignment while preserving expert oversight where interpretability and clinical judgment remain important.

The benchmarking results further support the effectiveness of this approach. The framework showed strong schema-level alignment and solid value-level performance across the major comparable clinical domains, with average results that remain strong for practical harmonization use. These findings suggest that the system is already well positioned to support scalable and analytically meaningful standardization of heterogeneous clinical datasets. Overall, the framework provides a practical route for transforming source EHR data into more consistent, reviewable, and OMOP-aligned representations, while retaining the transparency needed for trustworthy downstream use.

## Acknowledgments

The authors thank Krutika Gaonkar for their valuable feedback and support.

## Appendix

